# Fibroblast Growth Factor-induced lens fiber cell elongation is driven by the stepwise activity of Rho and Rac

**DOI:** 10.1101/2023.12.03.569812

**Authors:** Yuki Sugiyama, Daniel A. Reed, David Herrmann, Frank J. Lovicu, Michael L. Robinson, Paul Timpson, Ichiro Masai

## Abstract

The spheroidal shape of the eye lens is critical for precise light focusing onto the retina. This shape is determined by concentrically aligned, convexly elongated lens fiber cells along the anterior and posterior axis of the lens. Upon differentiation at the lens equator, the fiber cells increase in height as their apical and basal tips migrate towards the anterior and posterior poles, respectively. The forces driving this elongation and migration remain unclear. We found that membrane protrusions or lamellipodia are observed only in the maturing fibers undergoing cell curve conversion, indicating lamellipodium is not the primary driver of earlier fiber migration. We demonstrated that elevated levels of fibroblast growth factor (FGF) suppressed the extension of Rac-dependent protrusions, suggesting changes in the activity of FGF controling Rac activity, switching to lamellipodium-driven migration. Inhibitors of ROCK, myosin, and actin reduced the height of both early and later fibers, indicating elongation of these fibers relies on actomyosin contractility. Consistently, active RhoA was detected throughout these fibers. Given that FGF promotes fiber elongation, we propose it to do so through regulation of Rho activity.

**Summary statement:** FGF differentially regulates Rho- and Rac-activity at the lens equator to promote lens fiber cell elongation and migration.

## Introduction

Organ morphogenesis involves tissue elongation, invagination/folding, and branching; processes that are driven by cell division, cell shape changes, and cellular rearrangements (Lemke and Nelson, 2021). A combination of these events results in the formation of the spheroidal eye lens that is important to focus images onto the retina. The mature lens is primarily comprised of highly elongated convex fiber cells that are aligned with one another along their anterior-posterior axis (Fig. 1A). Lens growth consists of several discrete steps. The epithelial cells covering the anterior half of the lens proliferate and their progeny are gradually displaced posteriorly towards the lens equator. As they approach the lens equator, the epithelial cells exit from the cell cycle and begin the process of fiber cell differentiation. This differentiation process is driven by the elongation of the cells as their apical and basal tips migrate towards the anterior and posterior lens poles, respectively (see Fig. 1A). At the poles, the tip of one elongated fiber cell meets an opposing fiber cell tip to form a suture plane. Lens epithelial cell proliferation and fiber differentiation are thought to be regulated by changes in fibroblast growth factor (FGF) availability in the ocular media; the aqueous humor, that bathes the anterior half of the lens, contains a lower level of FGF sufficient to stimulate lens epithelial cell proliferation, while a higher level of FGF in the vitreous, that immerses the posterior half of the lens, induces and maintains lens fiber differentiation (McAvoy et al., 1991; Lovicu and McAvoy, 2005; McAvoy et al., 2017). While this mode of growth is widely accepted, given it explains many of the well-known histological features of the lens, fundamental questions regarding the nature of the forces that drive fiber tip migration and cell elongation remain unanswered.

**Fig. 1.**
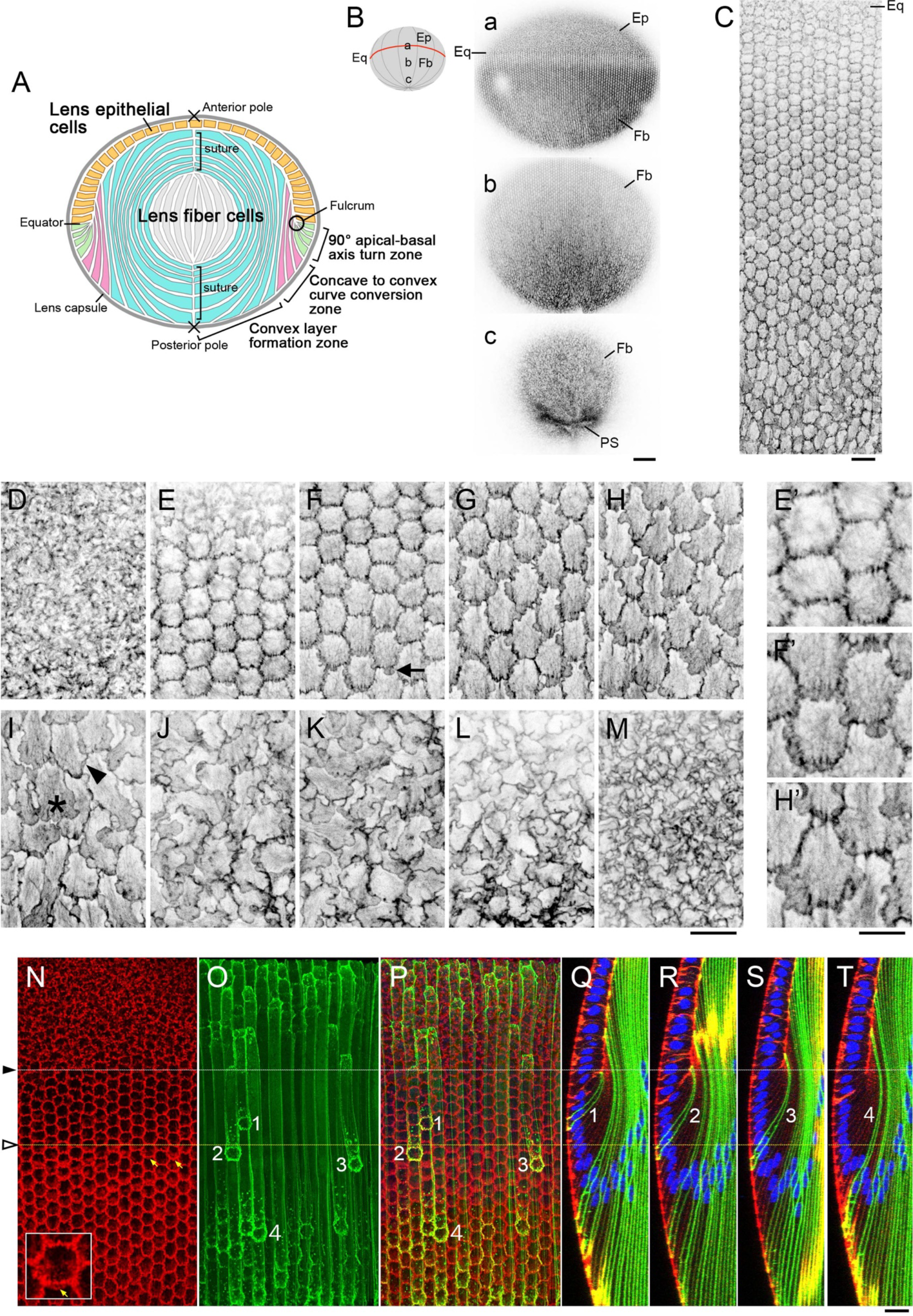
Morphological changes to the basal membrane of lens fibers during differentiation and maturation. (A) The lens of the mammalian eye includes an epithelial cell monolayer (yellow) and fiber cells (green, pink, blue, and grey). The innermost convex layers form around the older primary lens fiber cells that developed at early stages of lens morphogenesis (grey). (B) A whole lens isolated from a six-week-old wild-type mouse stained with phalloidin and visualized using confocal microscopy to show the distribution of actin filaments on the basal cell membranes. Sequential images were obtained starting from the equator (a) and extending to the posterior pole (c). (C) Morphological changes to the basal tips of fibers from a four-week-old lens during their differentiation at the lens equator. The posterior pole is at the bottom. (D-M) Higher power views of the basal membrane of fibers in lens tissue from a nine-week-old wild-type mouse. Magnified images of E, F, and H are shown in E’, F’ and H, respectively. The arrow in F indicates a small membrane protrusion extended to the posterior side of a slightly elongated hexagonal basal membrane. In I, the asterisk shows a basal tip with a fully developed membrane protrusion, and the arrowhead indicates the basal membrane with a regressed front. (N-T) A whole lens from a four-week-old mTmG x MLR39 Cre-transgenic mouse that expresses EGFP (green) in differentiating fiber cells, was stained with phalloidin (actin, red) and Hoechst 33342 (nucleus, blue). Yellow arrows indicate the earliest protrusions detected at the basal membranes of lens fiber cells (N, an enlarged image is shown in the inset). The basal tips of the lens fiber cells were numbered based on their respective locations from the youngest (1) to the oldest (4, O, P). (Q-T) Z-sections of lens fibers numbered 1–4 as shown in O and P). Fiber 2 had just detached from the fulcrum and had a broad apical tip rather than a pointed, tapered tip as seen in Fiber 1. Abbreviations: epithelial cells (Ep), Fiber cells (Fb), equator (Eq) and Posterior sutures (PS). Scale bars: B 100 µm, C 20 µm, D-M, 20 µm; E’, F’, H’, 10 µm. N-T 20 µm.

With closer evaluation of the fiber differentiation process, we note that the elongating fibers are located in three distinct zones; (i) the convex layer formation zone (Fig. 1A, cells in blue) is the largest zone and constitutes the major functional body responsible for optical refraction in the lens. This zone contains highly elongated fiber cells that are aligned along the anterior-posterior axis of the lens. These cells develop from (ii) precursors in the axis turning zone where the apical-basal axis is initially aligned perpendicular to the anterior-posterior axis at the lens equator (Fig. 1A, cells in green). Thus, the newly differentiated fiber cells undergo a 90° axis turn that is accomplished by the anchoring of their apical tips at the equatorial lens fulcrum. During this 90° turn, the differentiating fiber cells first develop their concave curve. After they accomplish this axis turn, in the following zone they convert from a concave to a (iii) convex configuration that we will refer to here as the curve conversion zone (Fig. 1A, cells in pink). For these cellular changes, we propose that multiple factors or signaling pathways drive the fiber cell movements, elongation, and curvature conversion.

Although the cortical fibers in the first two zones (the axis turn and curve conversion zones) do not contribute directly to lens refraction, their normal development is essential to facilitate the subsequent accurate formation of the convex fiber layer. Previously, Bassnett and colleagues examined the relationship between increased cell height and volume of the elongating fiber cells, reporting a proportional relationship between cell height and volume of the inner fibers, but no increase in the volume of the cortical fiber cells, concluding that the elongation of cortical fibers was primarily mediated by changes in cell shape and not cell volume (Bassnett and Winzenburger, 2003; Bassnett, 2005).

The equatorial fulcrum or modiolus is defined by the tapered apical tips of the differentiating lens fiber cells (Zampighi et al., 2000; Lo et al., 2000; Sugiyama et al., 2009). Formation of the fulcrum is disrupted in lenses from protein kinase C (PKC) λ or N-cadherin-deficient mice, that failed to form apical cell junctions (Sugiyama et al., 2009; Pontoriero et al., 2009; Logan et al., 2017). In lenses of these mutant mice, the newly differentiating fiber cells were prevented from undergoing an axis-turn and their apical tips were dislocated posteriorly as the lens fiber cells elongated. This indicated that while anchoring of the apical tips at the fulcrum is essential to support the axis turn and complete fiber elongation, other mechanisms may drive the increase in cell height in the absence of a functional fulcrum. Although the assembly of microtubule and actin filaments has been identified as a necessary feature of this process, the driving forces that promote their assembly still remain unknown (Audette et al., 2017).

Rho family GTPases are key regulators of actin dynamics and promote various cellular changes including cell shape and migration (Heasman and Ridley, 2008; Hall, 2012). In previous studies that examined lens tissue from mice devoid of either Rac1 or Abi2 [a component of the WASP-family verprolin-homologous proteins (WAVE) complex that is activated by Rac], the basal tips of lens fiber cells remained at or near the lens equator, and the concave curve exhibited by the newly-differentiated fiber cells did not switch to a convex configuration (Grove et al., 2004; Maddala et al., 2011). As Rac signaling drives cell migration by promoting the formation of membrane protrusions or lamellipodia at the migrating front, these results suggest that Rac-mediated basal fiber cell tip migration may be critical for the concave-to-convex conversion of lens fiber cells. While membrane protrusions at the basal tips of lens fiber cells have been documented using electron microscopy (Maddala et al., 2011), it is not clear whether these extensions are a specific feature of lens fibers undergoing curve conversion or generally associated with all lens fibers that are in the process of migrating from the equator to the posterior pole.

In this study, we first determined the lens zones where the membrane protrusions extended at the basal tips of lens fiber cells. We also examined how protrusion extension is controlled by FGF, the key inducer of lens fiber differentiation. We next examined the involvement of Rho-signaling for the elongation of lens fiber cells. In the current study, we applied inhibitors for Rho kinase and actomyosin to whole lens cultures and examined their effects on the cell height of fiber cells. To assess the activity of RhoA in the lens we also examined biosensor signals in lens fiber cells of genetically engineered RhoA Förster Resonance Energy Transfer (FRET) biosensor mice. We propose a new FGF and Rho/Rac-interactive network, whereby FGF promotes distinct steps of lens fiber morphogenesis through differential regulation of Rho- and Rac-signaling pathways.

## Results

### Posterior-directed lamellipodia are found specifically in fiber cells undergoing curve-conversion

To address whether Rac activity is required for curve-converting lens fibers, or generally associated with migrating lens fibers, we first examined the morphology of the basal tips of fiber cells from all lens regions of young to mature adult (3- to 9-week-old) wild-type mice. We isolated and stained whole lenses with phalloidin and generated surface views of fiber basal membranes using confocal microscopy. Whole lens staining revealed sequential changes to the basal membrane during fiber cell differentiation and cell migration from the equator to the posterior pole (Fig. 1B), particularly, the fiber basal tips that undergo dramatic morphological changes during fiber differentiation (Fig. 1C). When we examined the basal membrane morphology in greater detail, the basal membranes of the lens epithelial cells did not exhibit any distinctive morphology (Fig. 1D). In contrast, the basal membranes of the differentiating fiber cells transitioned to a regular hexagonal pattern (Fig. 1E). At this stage, the hexagonal basal membranes maintained discrete cell boundaries with no membrane protrusions detected (Fig. 1E’). Moving posteriorly, the hexagonal basal membranes appear to be stretched along the equatorial-posterior pole axis (Fig. 1F). Cells in this zone feature nascent membrane protrusions or lamellipodia that extend from the posterior sides of the fiber cell basal membranes (Fig. 1F arrow; Fig. 1F’). The number and size of these lamellipodia increased as the cells transitioned into the more posterior lens regions (Fig. 1G, H; Fig. 1H’), reaching a maximum (Fig. 1I, asterisk). Posterior to this point, the membrane protrusions diminished (Fig. 1I, arrowhead) and the basal membranes took on a packed appearance as the tips approached the posterior pole (Fig. 3J-L). At the posterior pole, the basal tips acquired a highly compressed pattern (Fig. 1M). Using whole-mount immunostaining of lens capsule, we detected Abi2 at the edges of the basal membrane lamellipodia, supporting that these protrusions were Rac-mediated structures (Fig. S1).

To specify the regions in which fiber cells extend these membrane protrusions, the basal surface images were linked to corresponding Z-section images to obtain zoning information (Fig. 1N-T). To facilitate tracking of the entire length of lens fiber cells, we used a transgenic mouse line (mTmG x MLR39-Cre) that expresses enhanced green fluorescent protein (EGFP) in axis-turning lens fiber cells. The mTmG mice express tdTomato, and Cre-mediated excision of the tdTomato allele induces EGFP expression (Muzumdar et al., 2007). Previously, we showed that MLR39-Cre expressed Cre in differentiating fiber cells (Zhao et al., 2004). EGFP was consistently expressed in the fibers undergoing curve conversion, but we found that its expression began randomly just before fiber differentiation or during the axis-turning stage (Fig. 1O,P). This sporadic expression pattern enabled us to precisely track the entire length of lens fibers.

We first confirmed that newly differentiated fiber cells that did not extend protrusions from the basal membranes were those with apical tips that were anchored to the equatorial fulcrum and were undergoing an axis-turn (Fig. 1Q). We then identified fibers with nascent membrane protrusions (Fig. 1N, arrows) with apical tips that had just recently detached from the fulcrum (Fig. 1R). This finding indicated that these fibers had completed the axis turn and had just initiated the process of curve conversion. The lens fiber cells continued to elongate and proceed with curve conversion at points beyond this boundary (Fig. 1S,T). This observation suggests that while curve conversion may be promoted by lamellipodia-driven active migration, as proposed earlier, the axis turn is induced by alternative mechanisms.

### FGF controls the capacity to extend membrane protrusions

To identify the mechanisms that control the behavior of differentiating fiber cells in each zone, lenses from adult mice were isolated and cultured for 24 h with inhibitors and/or growth factors. Lenses cultured in serum-free media lost the hexagonal shape in the basal fiber cell membranes, along with the generation of filopodia-like protrusions and stress fiber-like actin filaments in the axis-turn zone (Fig. 2A, B). Likewise within the curve conversion zone, the hexagonal profile of the basal membrane was deformed, accompanied by the extension of additional lamellipodia and an increase of the medial actin filaments (Fig. 2B). The excessive generation of actin filaments was effectively suppressed by the application of the Rho-kinase or ROCK inhibitor Y27632; however, the extension of the membrane protrusions was not suppressed (Fig. S2A-C). In contrast, the extension of these protrusions was suppressed in the presence of the Rac inhibitor NSC23766 (Fig. 2C). These findings suggest that while all lens fiber cells have the potential to extend membrane protrusions due to high levels of intrinsic Rac activity, this activity is suppressed in situ by unidentified mechanisms.

**Fig. 2.**
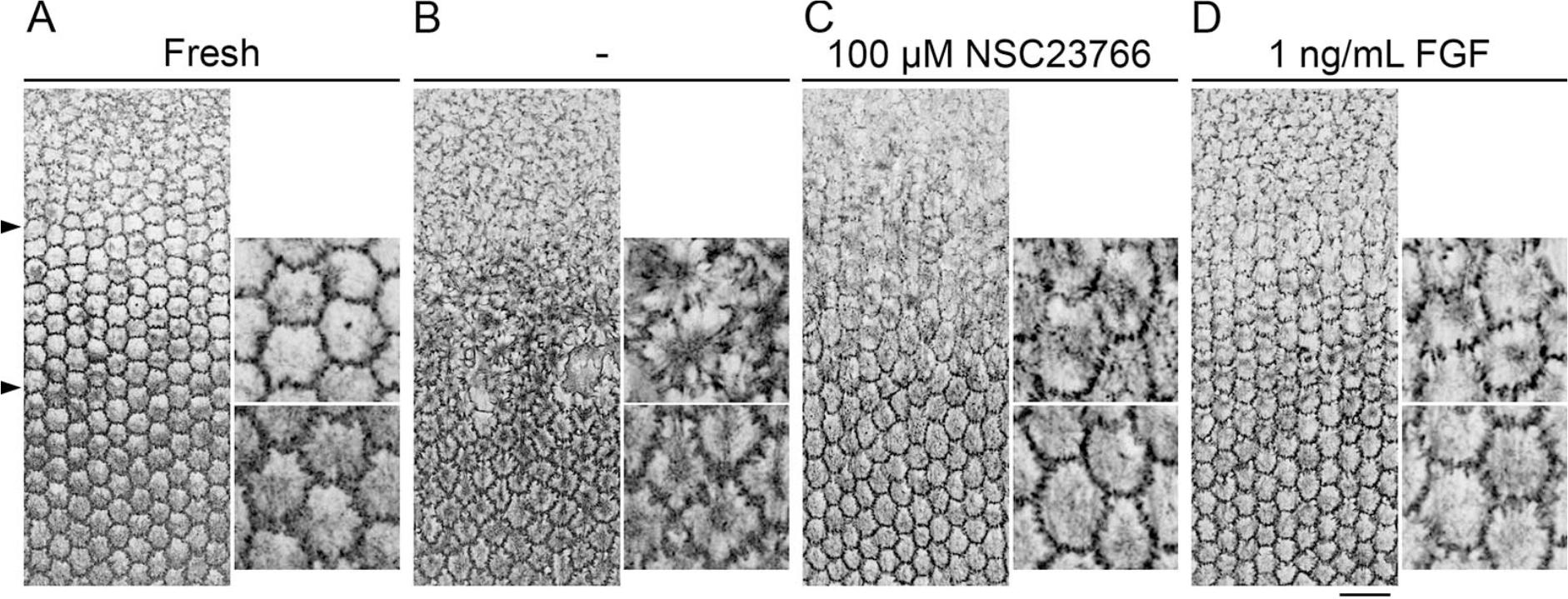
FGF-2 and the Rac-inhibitor suppress the constitutive extension of lens fiber cell basal membrane protrusions. Lenses from four-week-old wild-type mice were cultured in serum-free medium for 24 h and processed for phalloidin staining. The position of the fulcrum (upper arrowhead) and the boundary between axis turn and curve conversion zones (lower arrowhead) are indicated in (A); an image of a control lens that has not been cultured. Other images have been aligned at the fulcrum position, and enlargements of a portion of each zone are shown to the right of each panel. (B) After 24 h of culture, the hexagonal packing of lens fibers shown in (A) was deformed and spike-like membrane protrusions and medial actin fibers were detected. (C, D) The formation of spike-like protrusions was inhibited in cultures that included the Rac-inhibitor, NSC23766, and FGF-2. Scale bar: 25 µm.

To identify an endogenous factor that was capable of inhibiting Rac activity, we tested the role of FGF, the well-established inducer of lens fiber cell elongation and differentiation (McAvoy et al., 1991; Lovicu and McAvoy, 2005; McAvoy et al., 2017). When whole lenses were cultured with FGF-2 (1 ng/mL), it suppressed the formation of basal fiber membrane protrusions in both the axis turn and curve conversion zones (Fig. 2D). Interestingly, protrusion extension was suppressed more profoundly with higher concentrations of FGF-2; treatment of lenses with increasing FGF-2 doses (5 and 25 ng/mL) resulted in a more tightly packed basal membrane surface (Fig. S3). These results suggested that a FGF gradient may control the extension of membrane protrusions by modulating intrinsic Rac-activity. Thus, we propose a model in which relatively high FGF concentrations block the formation of protrusions in newly differentiated fibers. Subsequent reductions in FGF concentration permit the extension of lamellipodia only in the curve-converting zone.

### Inhibitors of actomyosin contraction reduce the height of cells in fibers

Although the observed posterior movement of the basal tips of fibers in the curve converting zone is most likely lamellipodia-driven, this cannot explain the anterior movement of the apical tips. It is also not clear how the basal tips of the newly differentiated fiber cells move posteriorly in the absence of membrane protrusions. One possibility is that the basal tips of fibers in the axis-turning zone are passively pulled in the posterior direction by the fibers with extended membrane protrusions in the curve conversion zone. Alternatively, Rho-mediated actomyosin contractility may drive an increase in cell height, thereby promoting fiber tip movement. Rho is a well-recognized regulator of actomyosin contractile activity, that plays key roles in the tail retraction of migrating cells, reorganization of cell junctions, and apical constriction during cell sheet folding (Heer and Martin, 2017). A reduction of the apical surface by constriction induces cell height increases and basal retention of the nucleus (Gelbart et al., 2012). We noticed that during axis turning, the nuclei remain adjacent to the basal membrane and move apically once they enter the curve conversion zone (Fig. 1Q–T). This suggests that the constriction is resolved once the fiber cells begin to undergo curve conversion. Thus, the observed increases in fiber cell height in the axis-turning zone may be the result of Rho-mediated actomyosin-dependent apical constriction.

To explore this possibility, isolated lenses were cultured with inhibitors of Rho-mediated contraction to evaluate their impact on fiber cell height within the axis-turning zone. In lenses cultured with or without DMSO (dimethylsulfoxide, inhibitor solvent) alone, axis-turning fibers showed minimal morphological changes compared to freshly-isolated lenses (Fig. 3A–C). In contrast, the addition of an inhibitor for ROCK had profound effects on the apical portion of the axis-turning fibers, causing abnormal space formation and disruption of the basal retention of the cell nuclei (Fig. 3D). Inhibitors of myosin (blebbistatin) and actin (cytochalasin D) also affected the position of cell nuclei in the axis-turning fibers (Fig. 3E, F). This response was not observed in lenses treated with NSC23766 (Fig. 3G), confirming that the defect was specific to actomyosin contractile inhibitors, and that it is not a Rac-mediated membrane protrusive activity.

**Fig. 3.**
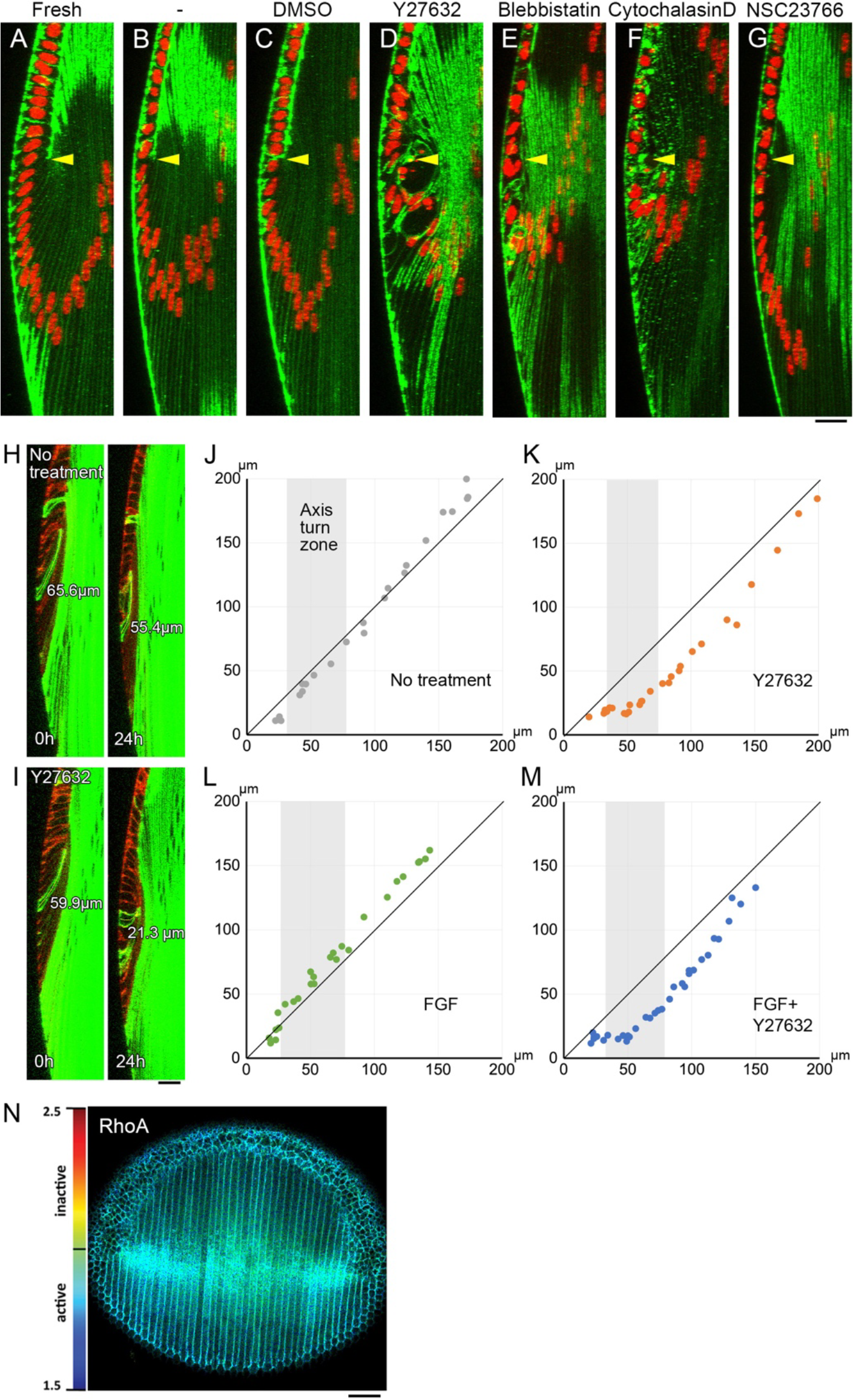
Inhibitors of cell contractility reduce cell height of newly-differentiated fiber cells. (A–G) Whole lenses from four-week-old wild-type mice were cultured with 1 ng/mL FGF and inhibitors of actomyosin contraction for 24 h, and stained with phalloidin (green) and Hoechst 33342 (red). Yellow arrowheads indicate the position of the equator. Control groups included (A) freshly-isolated lenses, (B) lenses cultured without any additions, and (C) lenses cultured in vehicle alone (0.2% DMSO). Inhibitors of Rho-associated kinases (50 µM Y27632, D), myosin (40 µM blebbistatin, E), and actin (0.25 µM cytochalasin-D, F) reduced the cell height of newly-differentiated fiber cells and induced a nuclear apical shift. (G) Rac inhibitor (100 µM NSC23766) did not replicate this response. (H, I) Whole lens isolated from an mTmG x MLR39 Cre mouse was cultured without (H) or with 50 µM Y27632 (I) and evaluated by live-image recording for 24 h. Changes in the height of individual fiber cells were tracked by EGFP expression (green). The tdTomato signal (red) was used to detect the location of the equator. (J–M) Fiber cell height (µm) was measured at 0 (x-axis) and 24 (y-axis) h. While slight reductions were observed in lenses cultured without FGF-2 (J), the addition of Y27632 alone resulted in substantial reductions in fiber cell height (K). Similarly, while the addition of FGF alone resulted in increased cell height (L), Y27632 induced a substantial reduction in the height of FGF-treated cells (M). (N) Representative FLIM images of RhoA. The conformational changes of the biosensors upon GTP loading to the RhoA motifs (i.e. activation of RhoA) result in a reduction in fluorescent lifetime of the donor fluorophore. In the lifetime maps high RhoA activity is represented by blue to green colors while yellow to red colors indicate low RhoA activity. Scale bars: A–G, 20 µm; H–I, 20 µm, N 50 µm.

We next assessed the change of cell height using live-image recordings of lenses from transgenic mice engineered to express EGFP in fiber cells. In these experiments, changes in the height of EGFP-positive single cells were tracked for 24 h (Fig. 3H–M). Our results revealed a subtle reduction of fiber cell height in lenses cultured without supplements (Fig. 3J). In contrast, the addition of Y27632 resulted in a substantial reduction in cell height (Fig. 3K). Furthermore, the addition of 1 ng/mL FGF-2 induced an increase in cell height (Fig. 3L); however, this response was blocked by the addition of Y27632 (Fig. 3M), indicating that the generation of the elongated cell shape of the axis-turning fibers relies on ROCK activity.

We noticed that the reduction of cell height induced by Y27632 was not only observed in the axis-turning but also in curve-converting fiber cells, suggesting the cell height increase of this zone is also regulated by Rho signaling (Fig. 3K, M). In support of this finding, we detected RhoA activity in both axis-turning and curve-converting fibers, using fluorescence lifetime imaging microscopy (FLIM) in a previously reported genetically engineered RhoA FRET biosensor mouse (Nobis et al., 2017, Fig. 3N). Taken together these observations suggest that fiber elongation in both axis-turn and curve-conversion zones is regulated by Rho-mediated actomyosin contractility.

## Discussion

The Rho family of small GTPases plays critical roles in diverse cellular events such as cell shape change and migration by modulating actin dynamics (Heasman and Ridley, 2008; Hall, 2012). In this study, we demonstrated that the morphological changes to the basal tips of lens fiber cells in the early two differentiation zones are regulated by ROCK and Rac. As discussed later, we propose a model in which the switching of Rho and Rac activity correlating to the distinct zones of fiber tip migration is regulated by different levels of FGF at the lens equator. Our findings also suggest that primary elongation of fibers is driven by ROCK-dependent actomyosin contractility.

### Different levels of FGF generate distinct Rac-active zones in cortical lens fibers

Results from previous studies performed in gene knockout mice revealed that Rac-signaling is a critical regulator of curve conversion in lens fiber cells (Grove et al., 2004; Maddala et al., 2011). While earlier results suggested that extension of lamellipodia promotes curve conversion, it was not yet clear whether this process contributes specifically to curve-converting cells or whether it was a general device used by lens fiber basal tips to promote cell migration to the posterior pole. Here, we showed that the extension of lamellipodia is a specific feature of fiber cells undergoing curve conversion and that extension of membrane protrusions is regionally controlled.

The results of experiments performed with our whole lens culture revealed that axis-turning lens fiber cells are intrinsically capable of extending membrane protrusions. It is likely that this intrinsic capacity to extend protrusions relied on the constitutively high level of Rac activity, since the formation of protrusions was suppressed by the addition of a Rac inhibitor. We also found that the addition of FGF prevented lamellipodia formation, speculating that *in situ*, FGF may normally suppress the formation of lamellipodia in fiber cells within the axis turning zone; however, a subsequent reduction in FGF levels may reactivate Rac at the start of curve conversion. Thus, we propose that at the basal tips of fibers that are exposed to the ocular media, different levels of FGF activity generate sequential Rac-dominant activity zones; relatively high levels of FGF in the axis-turn zone, negatively regulate Rac activity, and subsequent reduction in FGF activity observed in the successive curve conversion zone facilitates the reactivation of Rac and the development of fiber cell surface protrusions (Fig. 4).

**Fig. 4.**
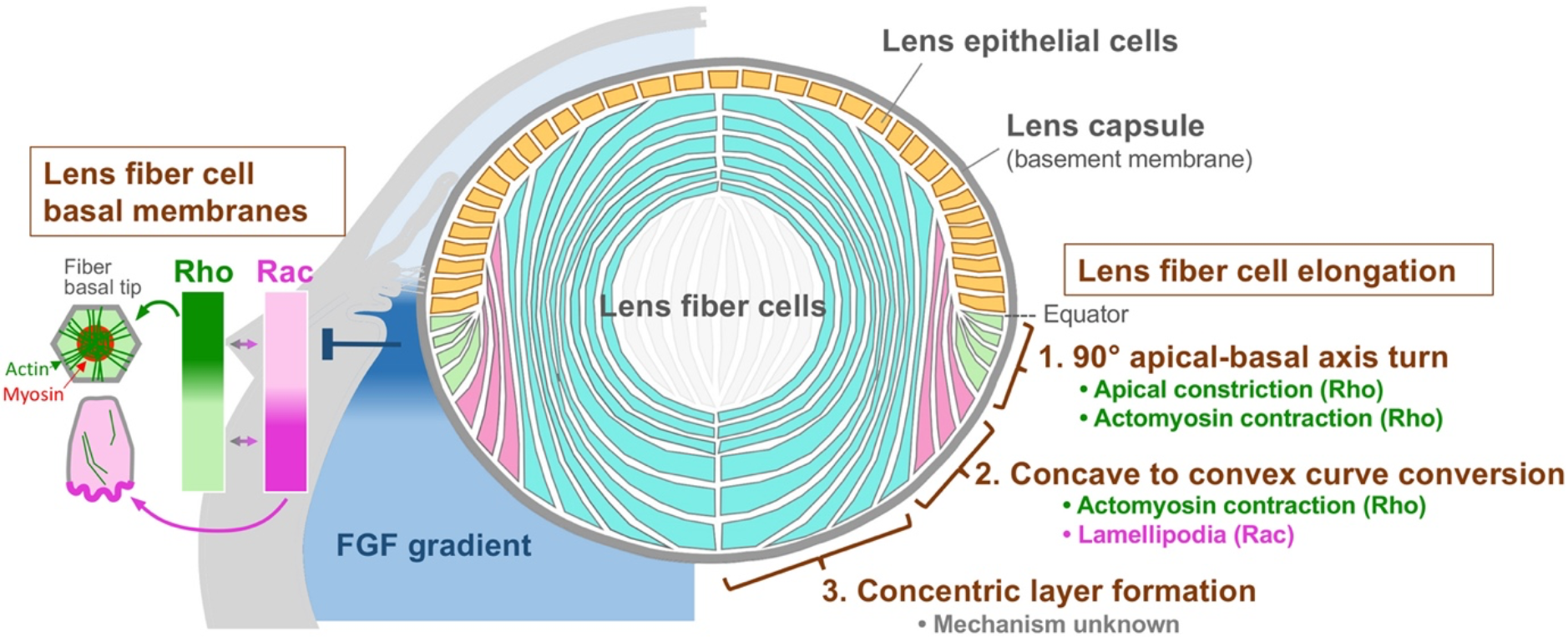
Model of lens fiber cell organization regulated by FGF. We propose a model in which lens fiber cells are divided into three zones based on their responsiveness to specific regulatory mechanisms. The elongation of the newly differentiated lens fiber cells in the axis turn zone (1) is regulated by Rho-mediated apical and lateral constriction. At the basal membrane, Rho promotes basal membrane complex formation. The basal tip migration of nascent fiber cells in the curve conversion zone (2) is mediated by Rac-dependent membrane protrusion or lamellipodia formation. The elongation of fibers in this zone is regulated by Rho-dependent actomyosin contraction. Further elongation of mature lens fiber cells in concentric layer zone (3) may depend on increases in cell volume mediated by an unknown mechanism. We predict that progression from zone (1) to zone (2) is controlled by changes in FGF concentration. Relatively high concentrations of FGF in zone (1) may suppress Rac-activity on the basal tips and thus prevent the formation of membrane protrusions, and reciprocally activate Rho to induce basal membrane complex. The FGF levels are slightly reduced in zone (2) and may serve to activate Rac and limit Rho activity, thereby inducing the extension of membrane protrusions.

Previous studies reported that progressive increases in FGF concentration stimulated proliferation, migration, and differentiation of lens epithelial cells (McAvoy and Chamberlain, 1989). Based on the comparatively lower FGF activity in the aqueous compared to the higher levels found in the vitreous (Schulz et al., 1993), it was proposed that lens epithelial cells bathed by the aqueous proliferate and migrate, and at the lens equator subsequently differentiate into lens fiber cells as they are exposed to higher FGF-activity of the vitreous (Lovicu and McAvoy, 2005). Our current results further suggest that, in addition to this antero-posterior FGF gradient, there may be a ‘hot spot’ at the equator that maintains higher FGF activity that diminishes slightly in the posterior region (Fig. 4). This specific configuration may allow lens fiber cells to develop membrane protrusions upon reaching the curve conversion zone. This transient elevated FGF signal may result from localized secretion by the ciliary body and/or the retinal marginal zone adjacent to the lens equator, consistent with earlier immunostaining experiments for FGF in these ocular regions (de Iongh and McAvoy, 1992).

Moreover, the expression of FGF antagonists, such as Sef, or MAPK/ERK antagonists Sprouty and Spred, may selectively negatively regulate FGFR activity to create this hot spot of FGF at the equator (Newitt et al., 2010; Shin et al., 2015; Wazin and Lovicu, 2020). Consistent with the higher FGF activity reported at this locale, downstream effectors of FGF signaling, including MAPK/ERK1/2 and ETS family transcription factors have been detected specifically at the equator (Lovicu and McAvoy, 2001; Upadhya et al., 2013; Xie et al., 2016). Taken together, these observations support our model for a heightened FGF activity at the lens equator.

We characterized the polarized extension of lamellipodia at the posterior side of the basal tips of curve-converting fiber cells. Previously, we showed that cortical fiber cells, that correspond to the curve-converting fiber cells featured in this study, exhibit planar cell polarity (PCP), with primary cilia on the anterior side of the apical membrane (Sugiyama et al., 2010). PCP is a mechanism that is regulated by Wnt-Frizzled signaling to arrange the cell axis within the plane of tissue (Butler and Wallingford, 2017). We propose that PCP is a characteristic feature of the curve-converting fiber cells that regulates both the direction of lamellipodial extension on the basal membranes and the polarized localization of primary cilia on the apical membranes.

### ROCK-mediated actomyosin contraction promotes fiber cell elongation

The current study revealed that lamellipodium-dependent migration cannot be the driving force underlying axis-turning. We subsequently examined the role of Rho-signaling, given that the hallmarks of Rho-induced apical constriction, including the induction of a wedged-cell shape, an increase in cell height, and basal retention of the nucleus (Gelbart et al., 2012; Heer and Martin, 2017) were detected in the axis-turning lens fiber cells. We observed that exposure to inhibitors of ROCK, as well as myosin and actin polymerization, resulted in the defects of apical cell tips and a reduction in cell height of axis-turning fibers, suggesting that ROCK-mediated apical constriction controls axis-turning and cell elongation.

We first predicted that inhibition of ROCK or actomyosin could affect fibers only in the axis-turn zone, but it also reduced cell height in curve-converting zones. Moreover, FLIM imaging in FRET biosensor mice showed RhoA activity in lens fibers of both axis-turning and curve-converting regions. This suggests that in addition to the apical constriction, fiber elongation in these zones is regulated by Rho-mediated actomyosin contraction, most likely on the lateral membranes (Fig. 4).

As FGF promotes fiber cell elongation, FGF may directly or indirectly activate Rho to promote actomyosin contraction that induces the observed increase in fiber cell height. Indeed, the application of low levels of FGF2 to the whole lens cultures was sufficient to promote fiber cell elongation; however, the control cultures without FGF showed a minimal reduction of fiber cell height. We also did not observe any reduction of the fiber cell height over 24 h when the lens was cultured with an inhibitor of FGF receptor signaling (SU5402, data not shown), indicating that once fiber cells elongate, FGF and/or other factors in the ocular environment are not required to maintain cell height/length. FGF has been known to activate actomyosin contractility in development of several tissues (Clarke and Martin, 2021). For example, during hindgut formation in chick embryos, the posterior endoderm undergoes Rho-A-dependent actomyosin contraction regulated by a postero-anterior gradient of FGF within the endoderm (Nerurkar et al., 2019). Thus, regional control of actomyosin contractility by a FGF gradient may be a common mechanism in tissue morphogenesis.

We detected ROCK activity also at the basal membranes of axis-turning fibers, that appears to be suppressed under normal situations not to form stress fibers. At high magnification objective lens, we detected radial actin bundles on the hexagonal basal membranes of these axis-turning fibers in fresh lenses (Fig. S4). Previous studies by Bassnett et. al. showed that in chick lens fibers, the hexagonal basal membrane formed a contractile complex composed of radial actin bundles and a medial myosin plaque (Bassnett et al., 1999). The distal actin bundles terminate at N-cadherin foci on the side of hexagonal cell boundaries, linked to actin bundles of adjacent cells, generating intercellular contractile forces to form a hexagonal lattice in the basal tips of lens fibers (Bassnett et al., 1999). We noticed that the radial actin bundles of mouse lenses were still visible in the fibers with nascent lamellipodia, but gradually disappeared as lamellipodium extension became apparent (Fig. S4). It appears that Rho activity dominates at the basal membranes of the fibers in the axis-turn zone, but then diminishes when Rac activity becomes dominant in the curve-converting zone. This suggests that at the basal membrane of fibers, activation of Rac may repress Rho activity in the curve-conversion zone as these small GTPases frequently exhibit mutually antagonistic activity (Mayor and Etienne-Manneville, 2016). This may result in a switch from an actomyosin-dominant contractile zone to a lamellipodium-driven migration zone.

In conclusion, our findings suggest that FGF differentially controls both Rho and Rac activity to induce early morphological changes of lens fiber cells that contribute profoundly to our understanding of the formation of the spheroidal lens shape essential for its transparency and focusing ability.

## Materials and methods

### Animals

The animal study protocols and procedures have been approved by the Animal Ethics Committee of Okinawa Institute of Science and Technology Graduate University, Japan, the University of Sydney, and Garvan Institute, Australia, and were conducted in compliance with the Australian code of practice for care and use of animals for scientific purposes. Wild-type C57BL/6J mice were used in this study. As we did not detect any gender-based differences, both male and female mice were used equally in all studies. The mTmG mice were purchased from JAX (stock number 007676, Muzumdar et al., 2007). This transgenic line carries a membrane-targeted tdTomato (mT) gene flanked by lox-P sites. Cre-mediated deletion of the tdTomato sequence activates downstream expression of EGFP that is targeted to the cell membrane (mG). MLR39-Cre transgenic mice express Cre recombinase specifically in differentiating lens fiber cells (Zhao et al., 2004). PCR primers for EGFP (forward 5’-ACGTAAACGGCCACAAGTTC-3’ and reverse 5’-TTGTACTCCAGCTTGTGCCC-3’, 370 bp) and Cre (forward 5’-AGGTTCGTTCACTCATGGA and reverse 5’-TCGACCAGTTTAGTTACCC-3’, 235 bp) sequences were used to genotype the mTmG and MLR39-Cre alleles, respectively. The PCR program was optimized to 94 °C for 2 min; cycling at 94°C for 10 sec, 60°C for 30 sec, and 72°C for 60 sec (x 35); 72°C for 5 min. Generation of RhoA and Rac1 FRET-based biosensor mouse lines was described previously (Nobis et al., 2017).

### Whole lens staining with phalloidin

Freshly isolated and cultured lenses were fixed with 10% neutral buffered formalin for 3 h to overnight at room temperature (RT). After washing (5 min to 1 h) with PBS, lenses were incubated with phalloidin (Atto 594-phalloidin, Sigma-Aldrich, USA) or Alexa Fluor 633 phalloidin (Invitrogen), and Hoechst 33342, in 1% Triton X100 supplemented phosphate-buffered saline (PBS) overnight at RT, or for 3 nights at 4°C. After washing (or without washing), a lens was placed between two strips of sponge in a culture dish filled with PBS. The lens was covered with a coverslip and evaluated by upright confocal microscopy (Zeiss LSM 710). Alternatively, stained lenses were placed between two strips of sponge in a glass-bottom dish filled with PBS for evaluation using inverted confocal microscopy (Leica SP8). Super-resolution images were obtained by Leica SP8 with a 100 x objective lens at zoom 5 and pinhole set to AU 0.4, and deconvolution was applied using Hyugens software.

### Whole lens culture with inhibitors

Eyes were removed and collected in a 35 mm culture dish containing 3 mL of M199 medium (11825015, Gibco) supplemented with 2 mM L-glutamine (25030081, Gibco), 100 U/mL penicillin - 100 μg/mL streptomycin (15140122, Gibco), 2.5 μg/mL Amphotericin B (15290018, Gibco) and 0.1% bovine serum albumin (BSA, A7030, Sigma-Aldrich). Lenses were isolated and transferred to a fresh dish containing 1 mL of M199 with 1 ng/mL FGF and/or inhibitors. After 24 h of culture in a standard 5% CO_2_ chamber at 37°C, lenses were fixed and processed for phalloidin-staining as described above. Stock solutions of NSC23766 (Rac inhibitor 553502, Calbiochem) and Y-27632 (Rho-kinase/ROCK inhibitor 688000, Calbiochem) were prepared with deionized H_2_O. Stock solutions of (S)-nitro-blebbistatin (myosin inhibitor 13891, Cayman) and cytochalasin D (actin polymerization inhibitor 11330, Cayman) were prepared in DMSO. Recombinant human FGF basic/FGF-2 (233-FB-025, R & D Systems) was dissolved in 0.1% BSA in PBS. We first determined the concentrations of inhibitors that induced cytotoxicity, then used lower concentrations to investigate the effect on fiber cell elongation. The concentration that induced abnormal appearance of lens fiber cells when cultured with 1 ng/mL FGF were 100 µM (Y27632), 50 µM (blebbistatin) and 200 µM (NSC23766). Cytochalasin D altered the filamentous actin pattern at 0.1 µM but the anterior shift of the nuclei was only observed above 0.25 µM. All experiments were replicated independently at least 3 times for each of the inhibitors.

### Live-imaging of whole lens culture

A whole lens was held at an angle with the equatorial side down for the live-imaging procedure. An mTmG lens carrying the MLR39-Cre allele was placed in a small hole in the center of an agarose pad that was prepared by loading 1 mL of 2% agarose (SeaKem GTG, Lonza) in M199 on the surface of 35 mm glass-bottom dish (3970-035, IWAKI). Once the agarose set, 3 mL of M199 with or without 1 ng/mL FGF-2 was added to the dish. The lens culture was maintained at 37°C in a 5% CO_2_ atmosphere in a stage-top chamber during live-image recording. Y-27632 was added to the dish on the microscope stage immediately before recording. Lens fiber cell height was measured by manual tracing using ImageJ/Fiji software.

### RhoA FLIM-FRET imaging

Lenses of 6- or 16-week-old RhoA-FRET biosensor mice (Nobis et al., 2017) were isolated and placed in a hole of agarose pad on a glass-bottom dish as described above, filled with DMEM. Imaging was performed immediately as described previously (Nobis et al., 2017) using a Leica STELLARIS 8 DIVE multiphoton inverted microscope with a 25x 0.95 numerical aperture (NA) water immersion objective. A Ti:Sapphire femtosecond pulsed laser (Mai Tai eHP DeepSea, Spectra Physics) was used as the excitation source and tuned to 920nm for EGFP excitation (RhoA-FRET mouse). EGFP signals were detected by non-descanned detectors (505-535nm) in photon counting mode. Lifetime maps were generated using LAS X FLIM FCS software and a standard rainbow color look up table (LUT) was applied with 1.5 to 2.5 ns for EGFP (RhoA-FRET mouse). There was little to no variation between the three lenses observed.

## Acknowledgments

We would like to thank Dr. John McAvoy for providing us with the opportunity to commence this research and for his support throughout the project. We also would like to thank Dr. Koji Koizumi of Imaging Section (IMG) in the Okinawa Institute of Science and Technology Graduate University (OIST) for his assistance with the microscopic imaging experiments. The FLIM-FRET analysis was supported by the ACRF INCITe Centre (Garvan Institute, Australia).

