## Supplemental figure for "Fibroblast Growth Factor-induced lens fiber cell elongation is driven by the stepwise activity of Rho and Rac"

**Supplemental figures**

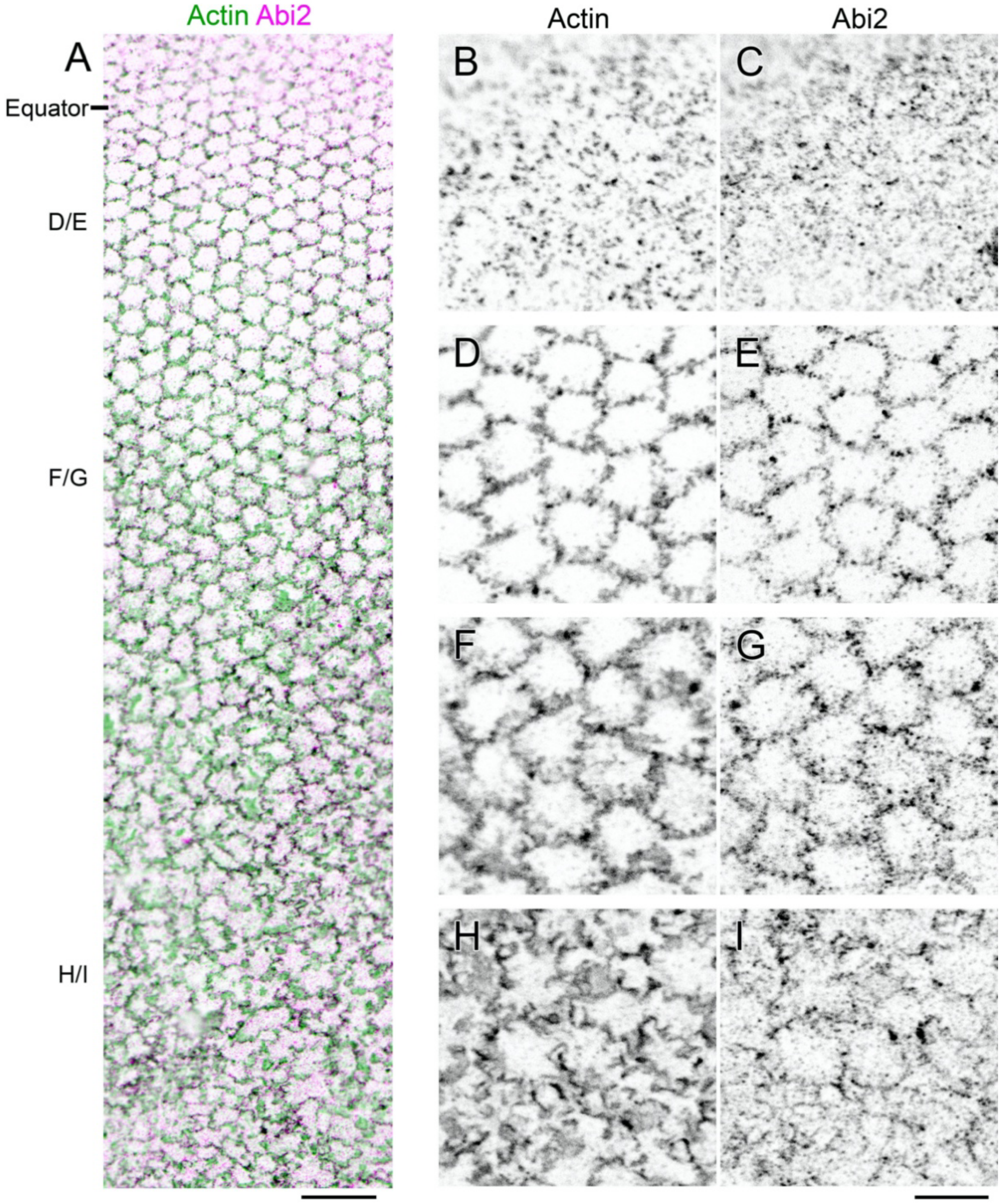


**Fig. S1. The membrane protrusions extended from the basal tips of lens fiber cells are immunostained with actin and Abi2 antibodies.** An isolated postnatal (P26) Wistar rat lens was fixed with methanol and the lens capsule was removed from the fiber mass. A whole mount was prepared with the basal tips of the fiber cells retained on the capsule and then processed for immunostaining (see Sugiyama et al., 2010). (A) A low power view of the wholemount posterior to the equatorial region. Anti-actin (green) and anti-Abi2 (purple) staining demarcated the cell boundaries. As shown, the hexagonal shape of the basal tips can be recognized up to a distance of 200 µm from the equator. After this point, the basal tips take on a more disordered appearance. (B-I) Detailed observations of the wholemount (A) immunolabeled with antibodies against actin (B, D, F, and H) and Abi2 (C, E, G, and I). The corresponding regions of panels D-I are indicated in A. (B, C) Actin and Abi2 were detected as individual spots within the lens epithelial cells. (D, E) Immediately after differentiation, both proteins were translocated to the hexagonal cell boundaries. (F, G) At the posterior end of the transitional zone, the cell boundaries stained with anti-actin and anti-Abi2 became slightly broader and less discrete. (H, I) As the hexagonal shapes underwent conversion to a more disordered appearance, both proteins were detected on emerging membrane protrusions. Actin was detected mainly at the edges and within the broad regions associated with the protrusions. Immunoreactive Abi2 was detected mainly at the edges of the membrane protrusions. Scale bars: A, 25 µm; B–I, 10 µm.

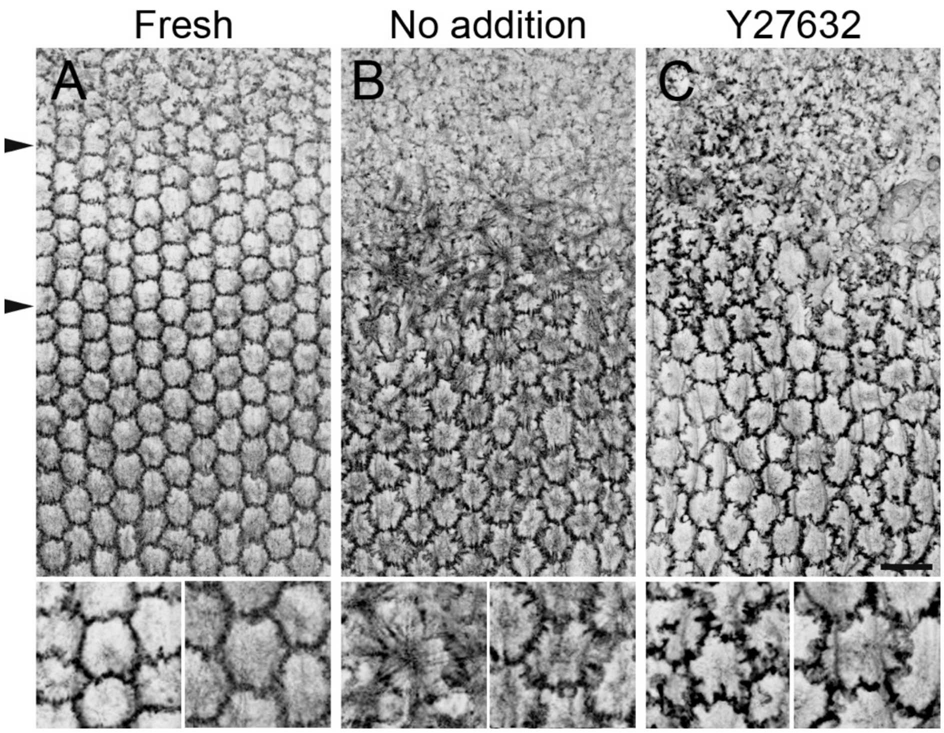


**Fig. S2. Inhibition of ROCK does not suppress intrinsic membrane protrusion formation on the basal membrane of fibers.** Four-week-old mouse lenses were cultured whole in the presence of 50 µM of the ROCK inhibitor Y27632 for 24 hours, similar to the experiments shown in Figure 2. Changes in the basal membrane morphology were assessed by phalloidin staining and observation using confocal microscopy. The ROCK inhibitor effectively suppressed the formation of stress fiber-like actin bundles but did not prevent the extension of membrane protrusions. Scale bar: 20µm.

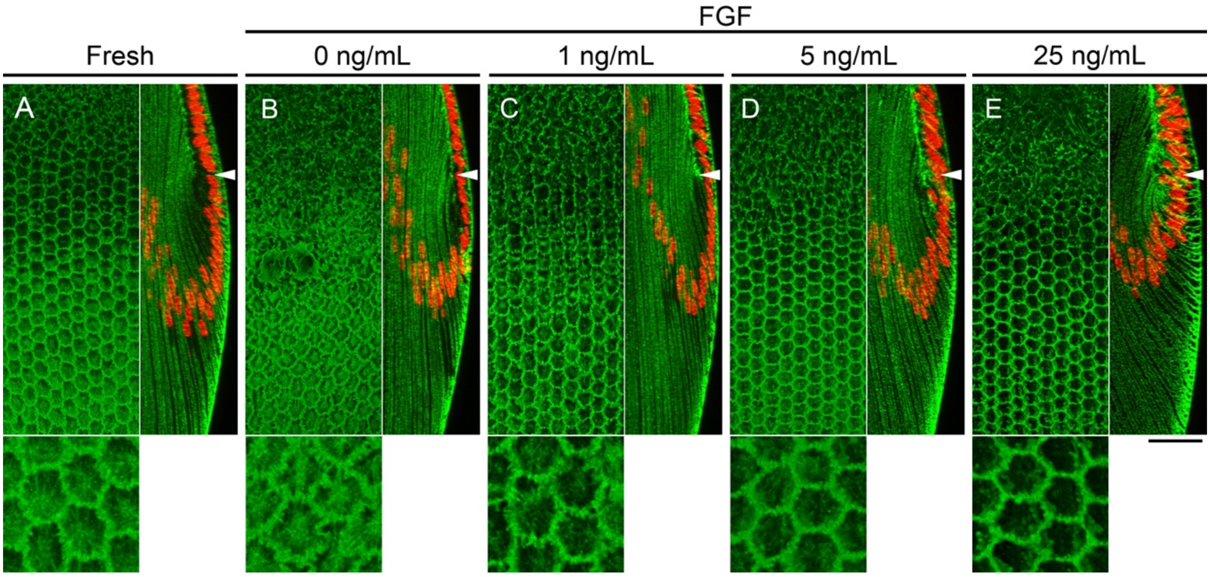


**Fig. S3. Higher concentrations of FGF result in reductions in the heights of fiber cells in the axis turning zone.** Whole mouse lenses were cultured with varying concentrations of FGF for one day and stained with phalloidin (green) and Hoechst 33342 (red). Surface three-dimensional projections (left) and reconstituted Z-sections (right) are shown. The equator or fulcrum positions are indicated by white arrows. Fresh (uncultured) lens tissue is shown in (A), and lenses cultured in serum-free medium without FGF-2 are shown in (B). (C) Although 1 ng/mL FGF-2 prevented thinning of lens epithelial cells and deformation of hexagonal basal membrane shape as shown in the non-FGF-treated lens (B), higher concentrations of FGF-2 (5 ng/mL, D; 25 ng/mL, E) resulted in tighter packing of the hexagonal basal membrane and the formation of accentuated concave curves. Scale bar: 40 µm.

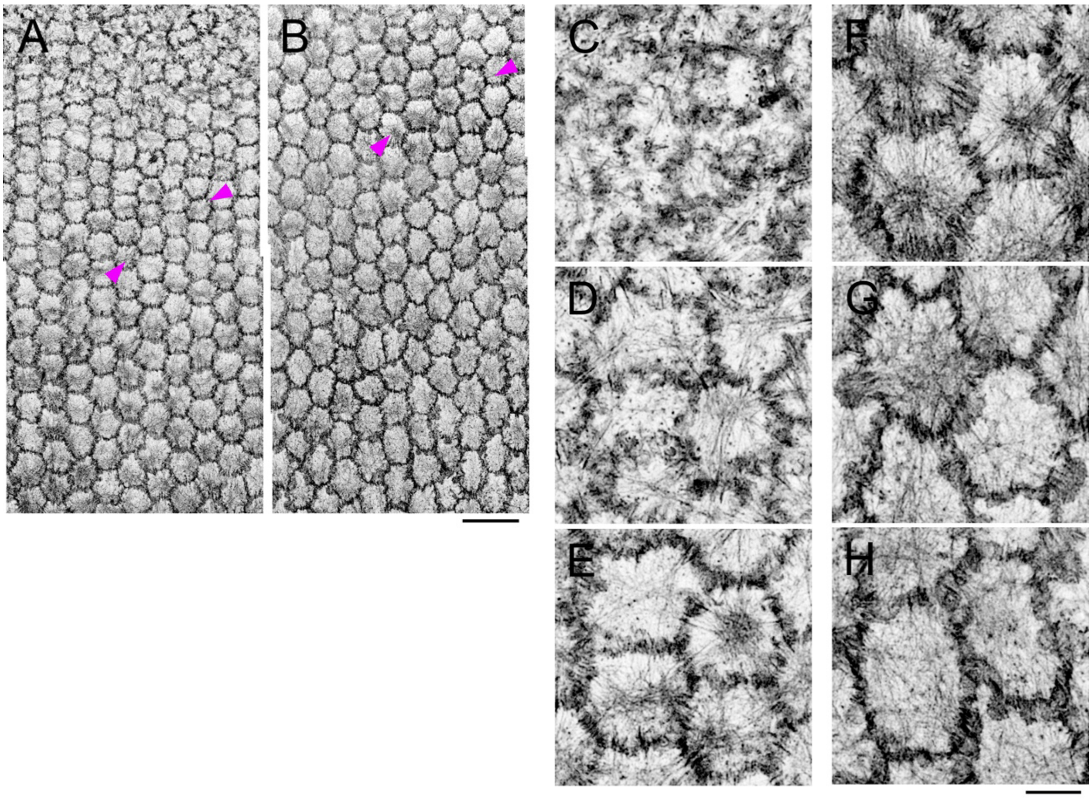


**Fig. S4. Dynamic changes of actin filament assembly on the basal membranes of lens fibers during maturation.** Confocal images of Phalloidin-stained whole lenses of postnatal day 15 mice imaged with a 63X objective lens for wider views (A,B), or a 100X objective with a 0.4 AU pinhole and deconvolution processing (C-H). (A,B) Basal tips of fibers undergoing differentiation to curve-conversion (A) and axis-turn to curve-conversion (B). Actin bundles occasionally showed inter-cellular alignment (between arrowheads). (C-H) A few actin filaments were seen in the basal membrane of the lens epithelial cells (C) but were arranged into radial bundles during fiber differentiation (D). In the hexagonal basal membranes, actin filaments occasionally formed a dense medial plaque and the radial bundles that emanated from it projected to the cell boundaries that appeared to be connected to the actin bundles of neighboring cells (E). The radial actin bundles were still visible in the fibers with nascent lamellipodia (F), but gradually disappeared as lamellipodium extension became apparent (G, H). Scale bars: A, B, 20 µm; C-H, 5 µm.
